# Probabilistic divergence of a TBM methodology from the ideal protocol

**DOI:** 10.1101/2020.07.05.160937

**Authors:** Ashish Runthala

## Abstract

Protein structural information is essential for the detailed mapping of a functional protein network. For a higher modelling accuracy and quicker implementation, template based algorithms have been extensively deployed and redefined. The methods only assess the predicted structure against its native state/template, and do not estimate the accuracy for each modelling step. A divergence measure is postulated to estimate the modelling accuracy against its theoretical optimal benchmark. By freezing the domain boundaries, the divergence measures are predicted for the most crucial steps of a modelling algorithm. To precisely refine the score using weighting constants, big data analysis could further be deployed.

## 1. Introduction

Topological detail of a protein provides valuable resource for the routinely deployed life -science studies including the annotation of disulphide connectivity pattern [1,2], functional annotation [3-5], molecular docking and cross-talk based studies [6,7], enzyme design [8,9] and drug development [10]. For technical and resource limitations, the structure determination methodologies have failed to bridge the ever-increasing sequence-structure gap of the protein sequences [11], and have started deploying the predicted models for building more accurate solutions [12].

Protein structure prediction has thus been the prime topic of research for several years. The widespread interest is not only because of the incompetency of structure determination methods to quickly solve the protein structures, but it is also attributable to numerous tribulations impeding our modelling accuracy. As per the worldwide test critical assessment of structure prediction (CASP), the modelling algorithms have been classified into free modelling (FM) and template-based modelling (TBM) which encompasses threading and comparative modelling protocols [13,14]. While the FM methods deploy the knowledge based potentials and physiochemical interactions and attempts to maximally search the conformational space possible for a protein sequence (target) for constructing its lowest-energy model [15], the TBM algorithms deploy the experimentally solved protein structures (*templates*) to construct a protein model. The TBM strategy is developed on basis of observation that highly similar protein sequences usually encode the same topology [16], and structural folds are robustly conserved across the functional diversifications [17,18]. It uses a reliable set of conserved structural folds to model a protein sequence [19,20]. Although, a probabilistically estimated threading methodology has recently attempted to estimate modelling accuracy for the selected templates and algorithmic constraints, it could not succeed for the absence of a robust assessment protocol [21]. A more accurate protocol to estimate the accuracy of a modelling protocol is still the need of the hour to consistently build more accurate protein models for all protein sequences.

## 2. TBM algorithm

An accurate TBM algorithm holds the key to consistently model a reliable protein structure for every target. However, against the available set of templates, its practical utility is dependent on the level of attainable accuracy of each of its steps. Although its every step holds its own significance to build an accurate model, a few steps viz. template search, template selection and combination, target-template alignment and model assessment are the most crucial ones to fix up the modelling accuracy, as has a lso been recently observed [22,23].

### 2.1 Template search

As a correct template set maximally spans the conformational space for a target sequence and serves as the basis to construct a near-native structure, it is crucial to significantly cover a target through functionally similar templates for building its near-native model [23]. Many template search matrices including the target-template sequence identity are not applicable in every modelling instance [24]. It is also suggested that some templates are prone to yielding false positive and erroneous matches for many target sequences, and thus create inconsistencies when a novel fold is being predicted with conserved sequence chunks [25,26]. The probabilistic estimation of the credibility of templates is still not sufficient to detect only the meaningful hits, as resulted for HHPred [27,28], and its threshold needs to be correctly defined to discriminate the functionally and topologically similar templates from the spurious hits. Although difficult to consistently screen the correct template(s), the profile-alignment based algorithms including PSI-BLAST [29], combinatorial extension algorithms [30], profile-HMMs [31], Jackhmmer [32] and HHblits [28] are usually deployed, and their threshold values for various scoring parameters have a great impact on the accuracy. A stringent and weaker template search threshold orderly fails to sensitively span the appropriate distant templates and exclude the distantly related and unreliable hits from the constructed profile [33]. Though CASP13 has extensively deployed the deep learning strategies over CASP12, the average GDT-TS scores of difficult/template-free target models is only found to increase to 65.7 from 52.9 (http://predictioncenter.org/casp13/). Hence, selecting good templates is the still the major challenging step to build highly accurate protein models.

### 2.2 Template selection and combination

For a relatively simple target with one/two continuous/discontinuous structural domains, multiple templates are easily available, and it makes the selection of a few evolutionarily closest biological hits highly complicated. On the other hand, when there is no significantly detectable sequence relationship among templates for a target, selection of correct templates again becomes a major limiting factor for the accuracy of the a TBM algorithm. Recently, CASP13 (http://predictioncenter.org/casp13/index.cgi), has orderly tested 13, 46 and 37 easy, single domain and unsplit domain targets sharing an almost complete coverage against multiple templates. The template selection has been shown to be especially difficult when such targets encode a novel fold with conserved substructures [26,34]. The easy targets include the single domain of a monomer or the compact domain of a multimeric structure. Although the model topology constructed through incorrect templates can never be energetically relaxed to the native structure [35], the topologically close models have been remarkably refined in CASP12 [36]. Hence, a correct selection of multiple templates, highly similar to each other for complementary augmentation, becomes essential to improve the overall target modelling accuracy [23]. The probability to predict such a set of correct, mutually complimentary and structurally similar templates becomes significant. Hence a good logically justified probability of such a crucial modelling step is needed.

### 2.3 Constructing a reliable target-template alignment

The modelling accuracy of a target is significantly dependent on the accuracy of its alignment against the template(s), which in turn depends on the proportion of correctly aligned and biologically significant residues. Profile-HMM-based algorithms deploy PFAM to avoid shifted alignment errors [31] and consider unaligned target chunks as single long insertions, and this hampers the overall modelling accuracy. As a reference guide, the structural information, inherent to the selected templates, has been also deployed by several protocols including combinatorial extension and HHPred [30,27]. However, most of these alignments need manual verification of the number, length and location of gaps, especially when a TBM-hard target is considered. The manually curated alignment becomes very useful to correct gaps incorporated in secondary structure elements, on the basis of sequence similarity and the secondary structure probability of the aligned residue chunks. Many modelling algorithms like MULTICOM [37,38] iteratively revisit the template selection and alignment steps to remove extraneous templates, or to overcome gaps localized in conserved regions.

A threading alignment usually attempts to be as accurate as a curated alignment, and through the maximum accuracy alignment algorithm, it has been shown to improve the accuracy of HMM-profile over the usual Viterbi scoring [39,28]. As the protein structures are robustly conserved over the sequence variations [40], many algorithms like MULTICOM [37,38] consider alignment ensemble to remove the irrelevant templates for overcoming the gaps bisecting the secondary structure/conserved regions. Hence, a probabilistic formulation is required to estimate the likelihood of a target-template alignment.

### 2.4 Model assessment

To select the most accurate model, the constructed/sampled decoy(s) are usually assessed against template/native structure on the basis of topological measures including root mean square deviation [41], template modeling score [42], global distance test score [43,44], contact area difference score [45], local distance difference test [46], SphereGrinder [47] and angular differences between successive atomic planes [48], and through solvent accessible surface area, hydrogen bonding network, pair-wise atomic interactions and molecular packing [49]. The global [50] and local [46,45] assessment scores are orderly used to select a topologically more accurate structure and find the inaccurately modelled regions for guiding the conformational sampling, usually deployed to reduce the non-physical atomic clashes [51]. However, even within an inadequately sampled landscape, multiple sampled models are usually close to the native topology [52,53], and for consistently selecting the most accurate model, a probabilistic estimation is needed to predict the likelihood of a protein model to be the correct near-native conformation.

Although a model-sampling-cum-assessment protocol has been developed through a probabilistic search [54], it could not prove to be successful. Although it shows that a soft-energetic bias steers the model sampling towards a diverse decoy dataset less prone to energetic artifact structures, it also demonstrates the algorithmic limitations of the usually deployed energy functions Hamiltonian and Rosetta. Recently in CASP13, for an incompetent assessment protocol, D-Haven could only construct a model with a GDT_TS score of 0.8293 for the target T0955 in contrast to the respective score of 0.9512 of the top-ranked structure predicted by MESHI’s lab (http://predictioncenter.org/casp13/).

As an attempt to deploy several diverse protocols more reliably, numerous meta-server algorithms have been developed for various modelling steps [55-57] as a hopeful attempt to deploy the majority of algorithms for building more accurate models. However, for consistently predicting the reliable models for all targets, the TBM algorithm still needs more accurate measures for evaluating the credibility of the modelling steps, in terms of the divergence score. It would empower our algorithms to quickly proofread the modelling strategy and rebuild an improved model. The further empowered TBM methodologies would provide credible models as the scaffolds to determine more accurate structures from the low-resolution experimental methodologies.

## 3. Probabilistic assessment of modelling steps

The standard practice of selecting the top-ranked templates is not consistently correct with a single unique algorithm. As TBM algorithms usually deploy a common set of independently implemented steps, it is possible to define an independent probabilistic assessment score for each step. Since a modelling strategy does not have an absolute assessment measure for every modelling step, we resort to estimating a measure of divergence for a modelling protocol from the results of computationally-feasible and the best-possible protocol with the highest modelling accuracy. Evaluating the modelling strategy of a predicted model this way against the one for the best possible theoretically feasible algorithmic structure, the credibility of a modelling methodology/step could be estimated. The score would be useful to find the most accurate protocol for each modelling step and improve the modelling accuracy.

### 3.1. Template search

It is well proven that the single-domain protein sequences, encoding utmost 200 residues, could have an average coverage of 70% against the templates and the predicted models could be within an RMSD of 5Å from their native state [58,18]. A target sequence usually shares a notable similarity against several templates/substructures, although the perfect set(s) of the non-redundant template(s) could maximally include a subset of N templates, including all the colocalized set of structural isoforms/topologically close structures. It is also logical to interpret that the likelihood of correctly screening such ideal entries is inversely proportional to the profile diversity [59].

Through profile-based strategies [27,60], integrated with the contact-map network [61] and deep-learning strategies [57], the templates are usually searched through the expanded sequence databases. An effective measure based on the colocalized alignment gaps, structural compactness and interactions with domain parsing need to be additionally assessed to evaluate the quality of an alignment for prioritizing the relevant templates only. As the effective template-scoring function/strategy is still missing, the protocol that screens the best set of templates as the top-ranked hits is actually the ideal one for a target sequence. While an ideal template is the experimentally determined target structure, the best set should be the union of all functionally related structures through which a near-native target model could be reliably constructed. However, the modelling methodologies fail to consistently screen this complete set, and as a tradeoff, heuristically fix a different set (K) to span the target to result into incorrect model topology. Hence, the logical divergence (Div_search_) for this step should be an absolute difference of the two sets and it should drastically increase when the additional/lesser templates are screened by a template-search algorithm, as formulated in the following equation I.

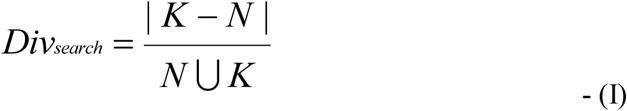

### 3.2. Template selection and combination

It clusters the screened templates as per the alignment location over the target sequence and selects minimal number of top-ranking and structurally complementary hits to gain the maximum coverage. An ideal template-set specifically differs in size and for entries. As each template-set, spanning a specific target segment, is independent of another group, the divergence measure of a modelling protocol can be reasonably approximated. If C_n_ builds the set of n groups each having a total of a_1_, a_2_, a_3_… a_n_ templates, and for every i^th^ cluster, k_i_ and n_i_ orderly defines the set of templates selected by a modelling protocol against the best possible set that could have built more accurate model, the logical divergence of this step (Div_selection_) can be approximated as equation II.

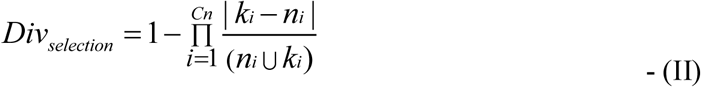

### 3.3. Constructing a reliable target-template alignment

For building a biologically meaningful target-template alignment and constructing a near-native target structure, the sequence similarity measure is usually deployed. However, when sequence similarity decreases, the evolutionary relationship among the related proteins cannot be reliably inferred [62,63]. Although the low-homology sequence alignments are curated through protocols like SFESA [64], the algorithm consistently constructing the most-optimal alignment is still not developed, i.e. the same protocol seldom builds the high accuracy model for the usual/CASP target sequence and its domains [23].

To estimate the divergence of the selected set of templates, a target-template alignment could be parsed into domains on the basis of Grishin plots [65] or DeepmetaPSICOV [66] or the alignment position against the number of gaps. As it could be accessed through a per-residue scoring difference for a common set of domains, the credibility of an alignment could be estimated for a template-dataset through the following equation, and it would be able to robustly assess even the partial accuracy of an alignment. If k_j_ and n_j_ orderly builds the set of templates selected by an algorithm against the ideal methodology for the j^th^ domain encoding l_j_ residues, and s_aci_ and s_opti_ is the score of the i^th^ residue for the j^th^ domain of the selected alignment against the ideal structural alignment, the divergence (Div_ali_) of a d-domain alignment can be estimated as:

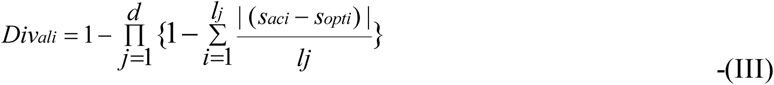

### 3.4. Model assessment

A global assessment of the constructed decoys helps to select the best model and guide the conformational sampling. It is still a major problem to select the most-accurate predicted model from the generated ones, especially if is a non-CASP target. All available measures are not consistently and unanimously correct to select the most accurate model for a target sequence. When domain orientation of a prediction is substantially variant than the closest template(s), measures like lDDT have proven to be more accurate, as recently observed for 11/65 targets in CASP13 (www.predictioncenter.org/CASP13/). However, for 43/65 oligomeric CASP13 targets, the assessment methodology is only found fairly competent for the monomer models and overall topology is not reliably predicted.

As has been repeatedly observed in all the CASPs, the best human predictions are found to be more accurate than the best TS servers which have only constructed structures of more consensus topology. The methodologies, building such an overall average topology for the top-ranked CASP server models, are not desirable for the advancement in the field. Despite the fact that for CASP13 target T0984, HHSearch could screen many templates with atleast 75% template-coverage, the best predicted model has only showed a GDT-TS and LDDT score of 57.94 (*monomeric*) and 64 (*oligomeric*). Furthermore, for these TBM-easy targets T0984-D1 and T0984-D2, the top-ranked models of KIAS-Gdansk and McGuffin has shown a GDT-TS score of 67.32 and 76.31 respectively, and it has again signified the requirement of a more robust assessment protocol.

Even when the models possess a global topological accuracy, the local stereochemistry should be correct and hence, the assessment team should work out at two levels. The conformational space of the models should be extensively sampled and it requires extensive data from both the modelling and assessment protocols. Therefore, for ignoring the near-native decoys (sd) and selecting the structurally divergent models from the constructed ones (cgd), the divergence of this step (Div_ma_) could be defined as

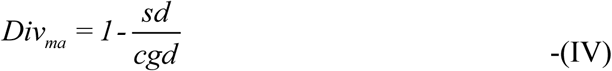

It has been recently quoted in CASP13 meeting that “Should the evaluation focus much more explicitly on the details of the model?” and it is implicated that the modelling protocol should primarily evaluate the topology, and if it is satisfactorily closer to the near-native conformation for almost the functionally important/conserved residues, the physicochemical details should be assessed.

As a linear scoring function fails to accurately assess the interdependency among the deployed sequence/structural parameters, the natural logarithm of the individual divergence scores should be considered to estimate the overall divergence of a modelling protocol. Considering α1-α4 as the four orderly defined normalization coefficients for the key parameters, the overall divergence can be postulated to be *Div*_tot_. Strategically refining our recently published modelling protocol [23] along with the divergence scores (*In progress*), with no extensive sampling, it shows a significantly higher modelling accuracy for a small randomly selected test-dataset of 5 CASP13 targets (Table1, http://predictioncenter.org/casp13/).

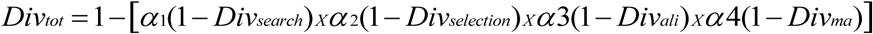

**Table 1:** Model assessment results for the 5 randomly selected CASP13 targets. A more accurate prediction is highlighted with boldcase letters. The developed strategy has shown a promisingly higher modelling accuracy than the best CASP13 models, and a negative score indicates superior CASP13 model.

| Target (length) | Top-ranked CASP13 model |  |  | Our model |  | Modelling accuracy improvement over the best CASP model |
| --- | --- | --- | --- | --- | --- | --- |
|  | Algorithm | Templates | TM-Score/ GDT-TS/ RMSD | Templates | TM-Score/GDT-TS/ RMSD |  |
| T0951_D1 (277) | IntFOLD5 | 3W04A, 5CBKA, 3W06A, 5DNUA, 1WOMA, 3QVMA | 0.971/93.284/ 0.952 | 5CBKA | **0.977/94.683/ 0.984** | 0.006/1.399/ 0.032 |
| T0961_D1 (505) | BAKER | 4Y9JB | 0.973/89.761/ 1.426 | 4Y9JA | 0.963/86.282/ 1.544 | -0.010/-3.479/ 0.118 |
| T0967_D1 (81) | BAKER-ROSETTA SERVER | 3W8G_B | 0.905/90.190/ 1.031 | 5HOKD | **0.910/90.506/ 0.980** | 0.005/0.316/ -0.051 |
| T0971 (186) | BAKER-ROSETTA SERVER | 4XBX_B | 0.972/95.769/ 0.729 | 3EBTA | **0.978/97.663/ 0.642** | 0.006/1.894/ -0.087 |
| T0976 (252) | DELClab | 2HHG_A, 1YT8_A | 0.687/41.224/ 3.681 | 1YT8A | **0.768/51.531/ 2.798** | 0.081/10.307/ -0.883 |
| T0976_D1 (120) | DELClab | 2HHG_A, 1YT8_A | 0.916/87.607/ 1.206 |  | 0.780/68.376/ 1.728 | -0.136/-19.231/ 0.522 |
| T0976_D2 (124) | A7D | N/A | 0.892/83.130/ 1.316 |  | 0.819/72.967/ 1.793 | -0.073/-10.163/ 0.477 |

## 3. Discussion

As observed in various studies [50,67], a more-robust assessment protocol seems a necessity for predicting the near-native protein structure. The earlier studies have sought to screen the topological/physicochemical errors in the predicted models for selecting the accurate structures [49,11,23]. To the best of our knowledge, no previous assessment protocol has probabilistically evaluated the modelling accuracy of a protocol. For estimating the accuracy of a modelling protocol, our postulated equations unanimously rank the complete modelling strategy and do not simply assess a lastly predicted model against its native state/template.

The modelling steps are analyzed under the given framework of quantifying internal consistency with respect to a scoring function. For instance, |K-N| and |N-K| variables within the template search equation define the number of excessive/trivial and omitted templates, as it has recently been shown that the accurate templates are often missed by even the top-ranked template search protocols [23]. Thus, the over-/under-selection will be appropriately dealt with this formula. Lastly, an equivalent assessment of all scoring terms is considered for an initial testing.

The best possible or most optimal algorithm is expected to deploy the best possible set of structures through the biologically meaningful alignment to yield the highest modelling accuracy, and in order that our analysis remains general, we compare this optimality at every step to another algorithm to evaluate the likelihood of its credibility. All assessments between procedures are made strictly under the premise of the same scoring function for both procedures, and without this assumption, there cannot be any kind of parity between the results of the two procedures. However, the divergence of a modelling methodology from the theoretical optimal protocol does not indicate the possibility of a more accurate algorithm. This is because the optimal procedure is computationally intractable for all practical purposes and therefore divergence of modelling results from the most optimal protocol is a measure to define the ranking methodology for each step so that the ill-performing steps could be curated. The lower the overall divergence score, closer is the prediction methodology to the ideally-best applicable algorithm. For the reason that some of the steps can affect the modelling accuracy much more significantly than the others, a well-trained set of different normalization coefficients is hereby required in the derived probabilistic equation. A major limitation of the method is that the coefficients (α1-α4) are strictly based on the deployed protocol. To add greater significance to these constants, big data analysis techniques such as machine learning should be further deployed to gather CASP information from the existing results of structure prediction algorithms and assign suitable values to the constants.

A combinatorial divergence score is parameterized to select the more accurate models and is logically shown to be more promising than the current assessment measures. Evaluating the modelling results for 5 randomly selected CASP13 targets against their native structures (*Parsed only for the officially assessed domain boundaries*), it indicates that our template-ranking strategy [23] has substantially improved the modelling accuracy. However, failure for some targets suggests the requirement of a threading protocol. Our article builds a chassis for developing a more accurate as well as automated modelling protocol that iteratively fixes a specific modelling step for each target sequence through the available sequence/structural constraints.

## Conclusion

As it is well understood that 51 tokens can be best delivered through two tokens: 50 and 1, and attempting any other combination is simply sub-optimal. We have described a standard decomposition of any protein structure prediction procedure and have constructed a divergence procedure for every modelling step. The proposed function works through comparison of a deployed strategy against the best theoretical protocol. As it is difficult to trace the algorithmic errors at every step of a modelling protocol, the strategy lays a logical chassis to develop a more accurate assessment measure that heuristically refine the modelling algorithm and consistently build a near-native model for every target.

## Acknowledgements

The author acknowledges the University and department for providing the required resources/support.

## In compliance with the ethical standards

### Ethical approval

This article does not contain any studies with human participants or animals performed by any of the authors.

## Availability of data and material

The constructed files/datasets analyzed in this study are available from the corresponding author on reasonable request.

